# SARS-CoV-2 Mediated Inhibition Of Respiratory Syncytial Virus

**DOI:** 10.1101/2024.09.02.610835

**Authors:** Elena M Thornhill, David Verhoeven

## Abstract

With circulation of SARS-CoV-2, fears about coinfection with other respiratory viruses such as influenza and RSV were significant, but the opposite was observed. Distancing/barriers played a major role in reducing other viral co-infections, however, some infrequent co-infections still occurred. We investigated the relationship between SARS-CoV-2 and RSV during coinfection to understand how they might compete or synergize. We found only RSV’s replication was significantly reduced when coinfected with SARS-CoV-2. Investigation of the mechanism revealed that the SARS-CoV-2 protein Nsp1 disrupts the RSV M2-2 protein but not the upstream M2-1 protein on the same biscistronic mRNA transcript. The impact of Nsp1 on M2-2 was not dependent on M2-2 being the second ORF in a bicistronic mRNA transcript, but likely from prevention of ribosomal termination-reinitiation necessary for M2-2 production. Additional viral ORFs from influenza A, influenza B, or Sendai virus dependent on the same or other ribosomal initiation mechanisms were tested and we found only influenza B M/M2 which likely uses a similar method as M2-2 was disrupted. Various M2-2 constructs, with/without the proposed site of ribosomal termination-reinitiating, co-transfected with Nsp1 and were in agreement that disruption to M2-2 expression occurs if the site of re-initiation was present upstream. These data not only suggest Sars-CoV-2 can outcompete RSV through suppression of M2, but may also point to potential ways to interfere with RSV by targeted therapies.

## Introduction

The SARS-CoV-2 virus was declared a pandemic by the WHO (World Health Organization) in March of 2020 and is responsible for approximately six million deaths thus far (Cucinotta and Vanelli, 2020). While vaccines have been developed, the rise of new variants undermines their effectiveness and, along with vaccine hesitancy or poor availability, has fostered the spread of the virus. Respiratory Syncytial Virus (RSV), like SARS-CoV-2, has the capacity for high case numbers. Moreover, in the U.S. alone, RSV is responsible for almost 60,000 hospitalizations of children under the age of five and ∼14,000 geriatric deaths each year. (Falsey and Walsh, 2005; Rose et al., 2018). Unlike SARS-CoV-2, RSV was first reported in the 1950s, yet our tools to treat RSV are seriously limited. RSV does not have an approved vaccine, and the approved prophylaxis treatment is expensive and only prescribed to premature (born prior to full-gestation) infants. While much recent attention has been given to SARS-CoV-2, RSV continues to threaten infants and the elderly with severe infections (Ducharme, 2021; Levine, 2022; McNiff, 2022).

Viral coinfections are a source of concern as the increased strain on the immune system of multiple concurrent infections can enhance disease severity in the infected individual (Nickbakhsh et al., 2019). Upon the arrival of RSV season in 2020, many in the northern hemisphere worried that RSV in tandem with SARS-CoV-2 would put an additional unbearable strain on the world’s health systems. In reality, RSV circulation numbers were lower than average (Olsen et al., 2021). While anti-COVID measures, such as masking, social distancing, and other viral limiting social strategies likely contributed to the decrease in RSV infections, there were still limited numbers of RSV infections. Any seasonal respiratory viruses would have had to compete with SARS-CoV-2 for hosts. While limited, there were instances of coinfections of other respiratory viruses and SARS-CoV-2, including RSV. However, RSV was not as prevalent as influenza in coinfections with SARS-CoV-2 (Hazra et al., 2020; Peci et al., 2021). RSV co-infections with SARS-CoV-2 has remained low or at least for the earlier SARS-CoV-2 variants (Halabi et al., 2022). The limited number of coinfections or infections in general seemed to indicate that RSV was struggling to compete with SARS-CoV-2. Then in the summer of 2021, RSV had an unexpected out-of-season outbreak (Ducharme, 2021). This outbreak corresponded to a drop in SARS-CoV-2 infections as the vaccination rates increased and more and more people were protected from SARS-CoV-2 (Ducharme, 2021; Olsen et al., 2021). The sudden shift to an out of season outbreak when SARS-CoV-2 rates were lower and a lack of typical RSV circulation during “peak” season suggested that these two viruses may have competed for host access. Thus, we sought to determine whether one virus might outcompete the other to access host cells.

Both SARS-CoV-2 and RSV are enveloped respiratory viruses that have RNA genomes. SARS-CoV-2 uses positive-sense RNA for its genome, while RSV uses negative-sense RNA. SARS-CoV-2 has the bigger genome at ∼30kB with 14 open reading frames and encoding ∼31 proteins (Accession number NC_045512.2). RSV has a 17kB genome with ten genes encoding 11 proteins (Accession number JX069799.1). Both viruses encode proteins that inhibit the immune and cellular response to the virus. Nsp1 is encoded at the 5’ end of the SARS-CoV-2 genome and is one of the first proteins produced in a polyprotein strategy (Brant et al., 2021). Nsp1’s primary function is immune response suppression and primarily inhibits the type 1 interferon pathway upstream and downstream of IRF3 (Schubert et al., 2020; Vazquez et al., 2021; Yuan et al., 2021). In addition to inhibiting interferon response, Nsp1 inhibits host mRNA translation by binding the 40S ribosomal subunit and inhibiting mRNA export from the nucleus (Lapointe et al., 2021; Zhang et al., 2021). Both Nsp1 and the host mRNA directly compete for access to the 40S subunit, which Nsp1 binds using its C terminal end in the mRNA binding groove during or directly after translation initiation (Lapointe et al., 2021; Schubert et al., 2020; Yuan et al., 2021). Residues K164 and H165 of Nsp1 are crucial for 40S binding, and amino acids 163-173 of Nsp1 are important for the translational suppression activities of the protein (Narayanan et al., 2008). Besides preventing 40S binding to mRNA, Nsp1 can induce cleavage or/and degradation of host mRNA and viral RNA, especially if the viral RNA contains certain IRES (internal ribosome entry sites) types. Moreover, cleavage or/and degradation of the RNA involves translational initiation factors and Xrn1 (an exonuclease), which Nsp1 can recruit (Yuan et al., 2021). Thus, dual infections may favor one virus over another through subversion of the host cell translational machinery or resource competition. There is evidence that coinfections, such as influenza and RSV, foster one virus over the other, but there are also coinfections where both viruses do not appear to influence each other (Esper et al., 2011; Marcos et al., 2011). SARS-CoV-2 appears to inhibit influenza replication, but little is known about the effects of the former on RSV coinfections. As Nsp1 can inhibit mRNA translation and other viral RNA, especially at sites of translation initiation, we wondered if Nsp1 could affect RSV translation.

RSV uses a start-stop mechanism to transcribe its mRNA. In the stop-start re-initiation model, the polymerase (L) must proceed step-wise, starting at the 3’ end of the genome, transcribing Ns1 before transcribing Ns2, and so on (Fearns and Collins, 1999a). If the polymerase fully disassociates from the genome at any point in the transcription process (except for the L gene region), it must return to the beginning of the genome and start with Ns1 again. This mechanism produces a gradient of mRNA transcripts such that there is a larger amount of the early genes like Ns1 being produced than that of the later genes such as the polymerase, though some recent papers contradict this model in favor of a non-gradient pattern of expression (Collins et al., 2013; Piedra et al., 2020). The M2 gene of RSV is transcribed as a bicistronic mRNA that is translated into separate proteins, M2-1, and M2-2. The current model for translation of both proteins is a mechanism called RNA termination-reinitiation, which is briefly described in Figure 3.1 (Gould and Easton, 2005; Gould and Easton, 2007). M2-1 is an essential RSV protein and functions to ensure full-length mRNA is made. The other protein produced on its bicistronic mRNA, M2-2, is often overlooked (Fearns and Collins, 1999b). While not considered essential, M2-2 functions to aid in the switch of the virus from a primarily transcriptive state of the polymerase to a primarily replicative state to prepare the virus for egress from the cell. Loss of M2-2 is severely attenuating, although with increased infection time, the virus can overcome the loss and is still capable of replicating its genome (Bermingham and Collins, 1999). Figure 1 illustrates the process for generating the downstream M2-2 off the M2 bicistronic mRNA transcript. There is an identified RNA secondary structure ∼150bp upstream from the M2-2 start site that fosters re-initiation in frame with the M2-2 gene start codon. Furthermore, 260bp upstream may also enhance the level of re-initiation (Ahmadian et al., 2000).

**Figure 1.**
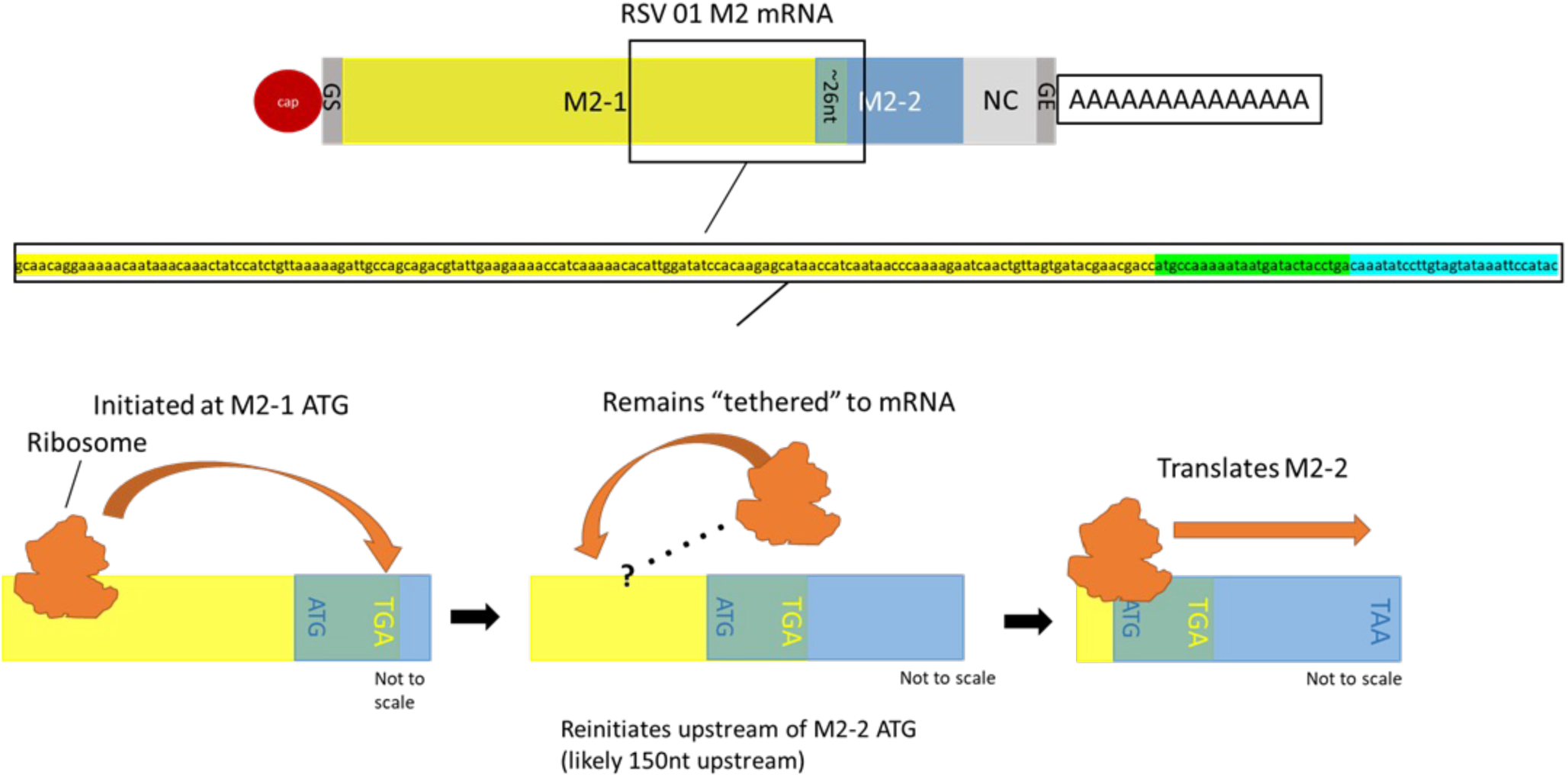
Mechanism of M2 translation through the termination re-initiation model of translation. Nucleotide sequence is specific to RSV 2001, but mechanism is universal to RSV strains.

We investigated the potential interaction between RSV and SARS-CoV-2 infection in permissive cells and showed that RSV infection is attenuated in the presence of SARS-CoV-2. Since Nsp1 has been recorded to interfere with the 40S ribosome, and the M2-1 to M2-2 switch involves the 40S performing a termination-reinitiation event, we hypothesized that the observed interference might be through Nsp1’s effect on M2-2 translation. Our data demonstrated that Nsp1 likely interferes with the 40S recruitment to the bicistronic mRNA nearby, at, upstream of the ATG start site of M2-2. Thus, our data suggest that SARS-CoV-2 interferes with RSV replication at least at the level of switching from transcription of mRNA to genome manufacture.

## Methods

### Viruses

RSVA 2001 was obtained from Beiresources (Manassas, VA) and expanded in Hep2 cells (ATCC, Manassas, VA), purified in PEG (polyethylene glycol) 8000 (Gias et al., 2008) and frozen in 3% sucrose in Dulbecco’s Modified Eagle Medium (Invitrogen, Carlsbad, CA). SARS-CoV-2 Wuhan strain was obtained from Beiresources and expanded in Vero E6 cells (Beiresources), tittered, and frozen in phosphate buffered saline (PBS, Invitrogen). All viral infections using whole viruses were performed in the ISU biosafety level III facility as approved by the ISU Institutional Biosafety Committee. Briefly, Vero E6 cells were infected with one virus or the other or in tandem at a multiplicity of infection (MOI) of 1 and allowed to infect for two hours before being removed and replaced with fresh media. Infections were incubated for three days post-infection, and the supernatant was removed and placed into Trizol reagent (Invitrogen) for RNA purification per the manufacturer’s suggested methodology.

### Plasmids

All RSV sequences are based on clinical RSVA 2001 strain. A pUC-T7-M2 plasmid was cloned (full M2-1/M2-2 bicistronic region under a T7 polymerase promoter), and site mutagenesis was performed using polymerase chain reaction (PCR) to add the 1xFlag tag and 6xHis tag. Additional pUC-T7 viral plasmids containing influenza B M/M2, Sendai C/P, and swine influenza H1N1 PB1/F2 were synthesized by Genscript (Piscataway, NJ). During their synthesis, a 1xFlag or 6xHis tag was added in-frame with the main or the secondary genes on the bicistronic plasmids. Nsp1 plasmid was made by RT-PCR from cDNA (Superscript III, Invitrogen) made from Wuhan genomic RNA (Beiresources), placing a T7 promoter sequence in front of the coding region and placing the sequence into pUC19. Additional Nsp1 mutations were made by site mutagenesis altering the R171 and K164 sites (or both) to A and E amino acids respectively. Transfections were done with 1-5μg of plasmids (most often 1μg) since the lower end consecutively gave a signal that was easy to stain and identify within cells that was consistent in each field of view. However, the T7 polymerase gene likely generated high levels of mRNA coding for the gene of interest (Piedra et al., 2020). Similar results (not shown) were obtained whether our plasmids had an ECMV IRES prior to the starting codons or not (Bochkov and Palmenberg, 2006) as the T7 polymerase system is highly efficient and the PRK antiviral system is known to be diminished in BSRT7 cells.

For our plasmids containing the M2-2 gene, we used 10 leading nucleotides prior to a kozak and ATG start site for M2-2 (called 10ATGM2 in text). We found this to be the most efficient expression of this protein using the internal flag tag for staining (data not shown). For the stated 150 or 260 nucleotides up-stream that contain the structure identified necessary for terminiation and reinitiation, we added a start ATG codon so that the primary protein product would be a truncated M2-1 gene but have the ability to make M2-2 with a flag tag if initiation reinitiation occurred (called 150 or 260M2 in the text).

For the influenza bicistronic PB1, Sendai bicistronic C/P, or influenza B bicistronic M2 genes, we synthesized these through Genscript placing a 6xhis tag in the upstream gene and flag tag (both in frame with their respective gene but out of frame but making irrelevant amino acids) in the downstream gene.

### Viral extractions and detections

SARS-CoV-2 N (nucleocapsid) protein qPCR system (IDT, Iowa City, IA) and RSV N qRT-PCR (Luna, NEB, Ipswich, MA) were detected under one-step qRT-PCR on a QuantStudio 3 (Applied Biosciences, Foster City, CA). Gene specific amplification and quantification was done similar to a prior study (Verhoeven et al., 2014) for some genes or individual RSV gene timing similar to another study (Piedra et al., 2020).

### Transfections

Cells were plated to ∼80% confluency before transfection. PCR products and plasmids were transfected into T7 expressing Baby Hamster Kidney Cells (22) (BHK: BSRT7) using transfections and antibodies as in the confocal section but using a Zoe fluorescent microscope (Bio-Rad, Hercules, CA). All transfections were carried out using X2 polymer (Mirus) using 1μg of plasmid where indicated or 5μg. Each was carried at using the manufacturer’s recommend ratios of plasmid to polymer and 30 minute co-incubation before adding to cells at 85% confluency. Transfections were allowed to go overnight. Our chose of 1μg for these transfections came down to that was the amount reproducible for each transfection and gave us good protein expression to see in our systems. Cells were then fixed using paraformaldehyde and permeabilized using a saponin buffer (Biolegend). Antibodies to 6x his or flag (Biolegend, Gentex) were incubated with cells in perm buffer for an hour before extensive washing and adding a secondary Alexa 488 or Alexa 594 for mouse or rabbit antibodies respectively. Cells were then washed again and then re-fixed.

### Transfection Imaging

All transfections were repeated at least three times in triplicate, five to ten fields of view were imaged for each transfection, and images shown are representative of the average field of view. An additional blinded reviewer confirmed the image results and counted the average numbers per field. Any contrast or brightness was applied equally across any matched image in all figures.

### Nsp1-RSV Co-transfection/infection

BSRT7 cells were transfected (similar as above) with pUC-T7-Nsp1 either 24hrs before, at the same time, or 15hrs after RSV (Long strain with mCherry reporter) infection at an MOI of 0.1. Cells were also infected with RSV alone, RSV with an empty plasmid (pUC19), or transfected with Nsp1 alone for controls.

### Statistical Analysis

Data was analyzed using Prism 8 software (San Diego, CA). Student T-tests were used on the data and p-values below 0.05 were considered significant.

## Results

### Coinfection Attenuates RSV

We first sought to determine whether one virus might be interfering with the other and performed co-infections in Vero E6 cells, which are highly permissive to both viruses. Specifically, we inoculated the Vero E6 cells at an equal viral MOI of 1. SARS-CoV-2 was either inoculated prior to (1hr before, not shown but similar to the other time-points), at the same time as, or after inoculation of the RSVA 2001 strain (3 hr after), and then changes in viral amounts were determined by qRT-PCR for both viruses using probes for the N gene. Of interest, the level of SARS-CoV-2 replication was equal regardless of when we put the virus into the cells suggesting that the viral replication was not interfered with by RSV. In contrast, we found a significant limitation on RSV replication when added at the same time or after (Fig. 2A) or before (not shown). These data suggest that SARS-CoV-2 infection interferes with RSV infection in some capacity.

**Figure 2.**
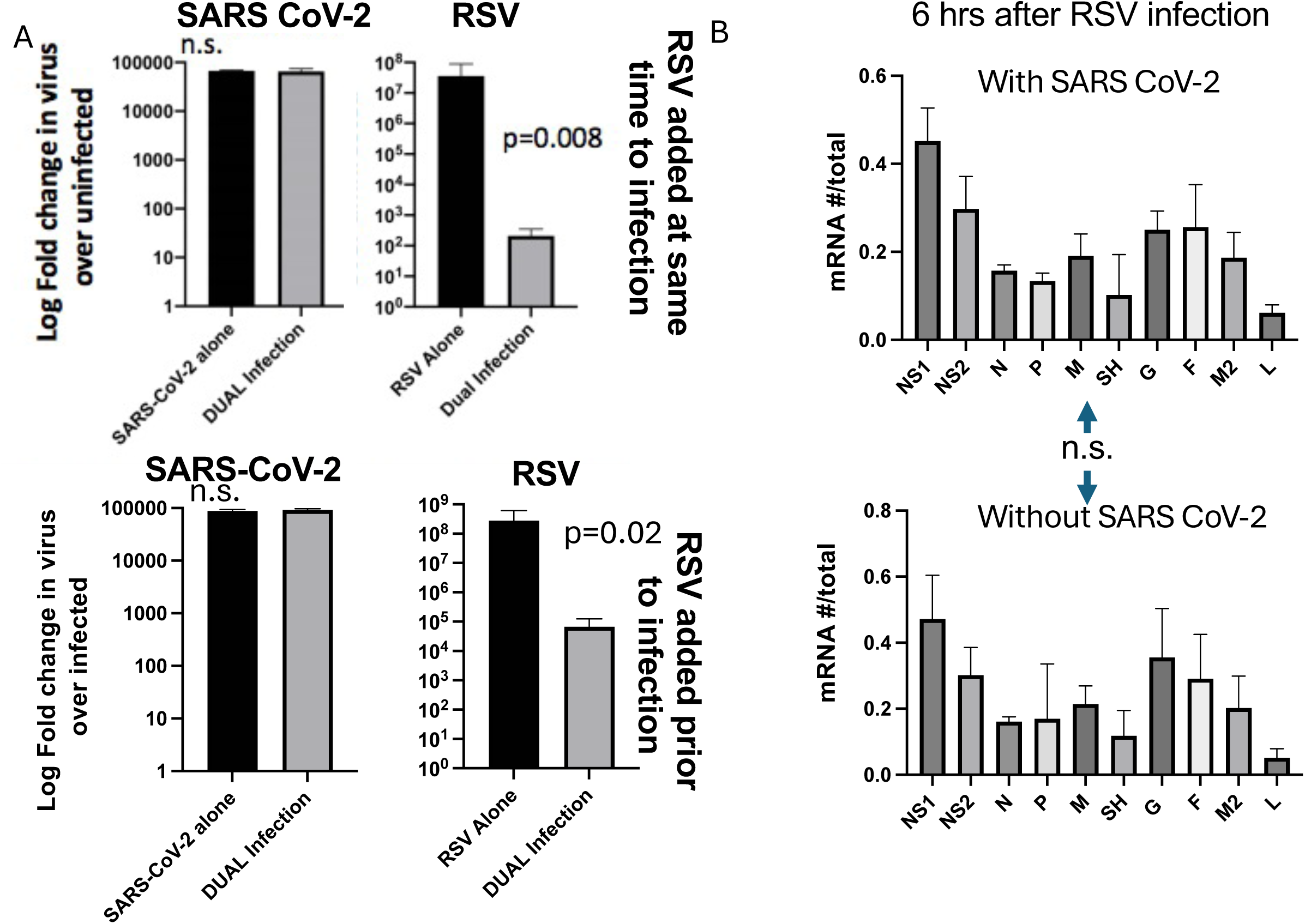
Co-infections between SARS-CoV-2 and RSV favors the former over the latter. **(A)** We infected SARS CoV-2 (Wuhan strain) either at the same time or after RSV 2001 strain (6 hrs). Both used an MOI of 1 into permissive Vero E6 cells. n=10 cell culture wells for two reps. Viral burdens were determined by qRT-PCR for each nucleoprotein. mRNA specific numbers from RSV were calculated from co-infected cells after SARS CoV-2 was added after RSV. n=3 using 3 replicates.

We next wondered if SARS-CoV-2 might block early replication in some capacity. We then infected RSV at the same time as SARS-CoV-2 and determined the relative expression of each gene 6 hrs after infection to determine if SARS-CoV-2 might be affecting RSV gene transcription (Fig 2B). Whether SARS-CoV-2 was added or not, there was no significant differences in the level of RSV gene expression. Similar results were obtained with RSV added after SARS-CoV-2. Thus, the data suggest that SARS-CoV-2 mediated interference is unlikely to be due to some effects on early RSV gene transcription.

### Nsp1 Inhibits M2-2 protein expression

The Nsp1 protein of SARS-CoV-2 is an inhibitory protein produced by the virus to limit host cell translation through interaction with the 40S subunit of the ribosome and interaction with mRNA transport mechanisms (Schubert et al., 2020; Zhang et al., 2021). Since M2-2 relies on 40S reinitiation (Fig 1), we speculated that Nsp1 might interfere with M2-2 production, thus limiting viral production. Thus, we next co-transfected plasmids coding for the SARS-CoV-2 Nsp1 protein and the RSV M2 mRNA (M2-1 and M2-2 proteins) into BSRT7 cells. Cells were transfected with Nsp1 and RSV M2 either alone or together in a 1:1 ratio were imaged for Alexa 488 (M2-1, 6xhis tag) or Alexa 594 (M2-2, Flag tag) and shown as individual channels (Fig.3A).

**Figure 3.**
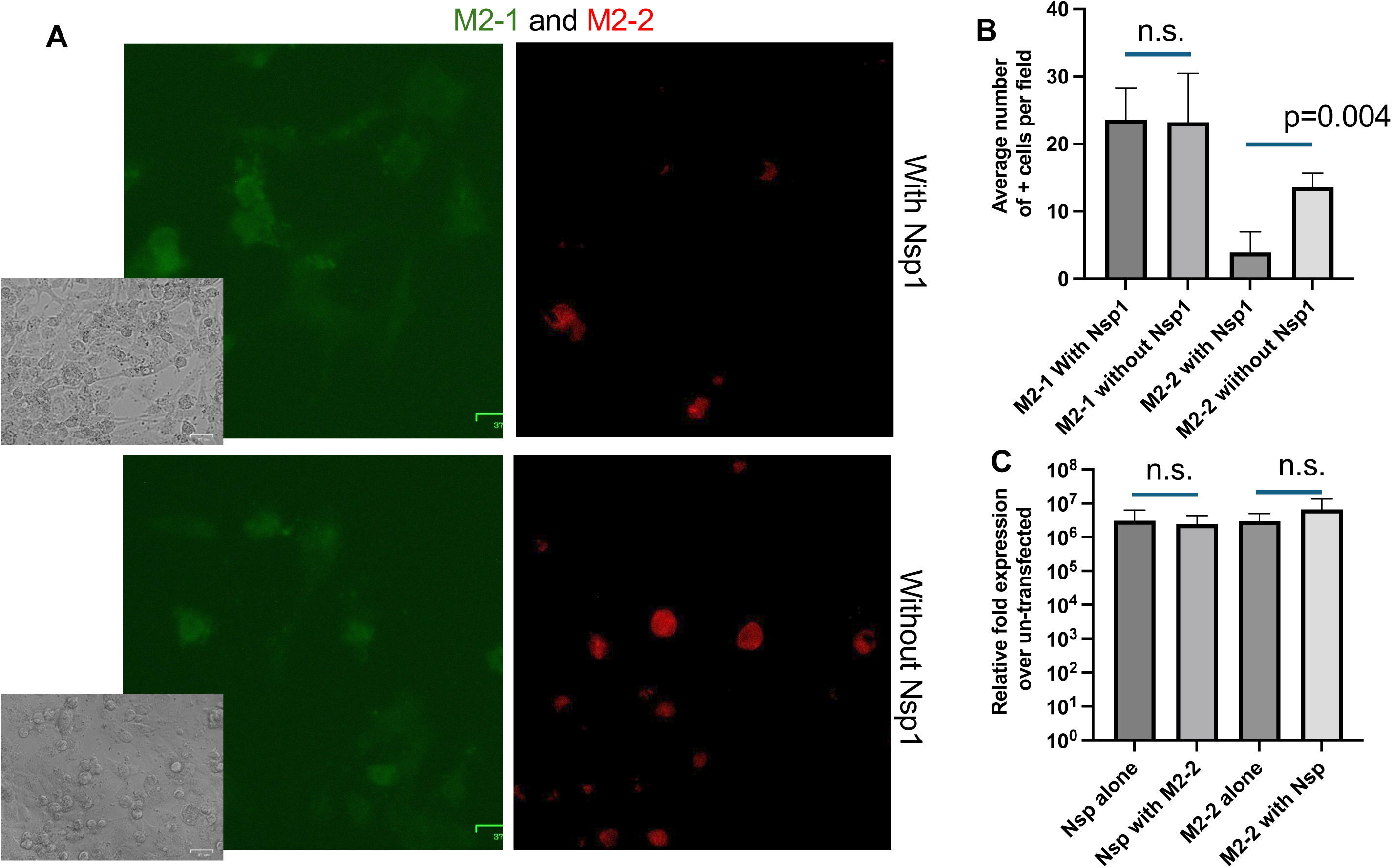
SARS CoV-2’s Nsp1 protein appears to interfere with M2-2 protein expression. Plasmids containing the Nsp1 gene after a T7 promoter or the full length M2 gene with a 6x his tag in frame with M2-1 and flag tag in frame with M2-2 were transfected into BSRT7 cells. Cells were dually intracellular stained for the Flag or 6xhis tag. **(A)** M2-2 alone or **(B)** with Nsp1 expression for M2-1 (Green, 6xhis) or M2-2 (red, flag) are shown. To ensure that our polymer transfection material was just not able to bind up more plasmid and deliver it in M2-2 alone transfections, we added a T7 promoter empty pUC19 control plasmid at the same amount as Nsp1 in the dually transfected cells. n=3 using 3 replicates Brightfield images are shown as insets showing 85-95% confluent cells for each image.

Within the average field of view of cells transfected with M2 alone, M2-1 showed expression in ∼17-30 cells within the field of view (Fig. 3B). M2-2 expression averaged ∼11-16 cells per field of view which is not surprising since M2-2 is made by termination and reintiation. (Figure 3B). While cells expressing M2-2 were lower than those expressing M2-1, M2-2 was detected throughout multiple fields of view when M2 was expressed alone in cells. In contrast, while M2-1 expression was similar whether Nsp1 was transfected or not, M2-2 expression differed in cell numbers expressing the protein when transfected with both M2 and Nsp1. M2-2 was not detected in some fields of view within these dual transfected cells and up to 8-9 cells per field in others.

To ensure no effects on transcript expression (and diverging from fig 2B) that might be affecting the outcome on both Nsp1 and M2 mRNA expression, we determined the relative expression by qRT-PCR (Fig.3C). There was no difference again in whether one transcript was present or the other suggesting again that M2-2 suppression is on the protein level rather than the transcript level.

Nsp1 severely knocked down M2-1 expression at a 5:1 ratio of Nsp1 to M2 (∼80% of cells per field of view to an average of ∼ two cells per field of view), indicating that Nsp1 is indeed capable of inhibiting M2-1 but not to the same effect as it does to M2-2 (data not shown). For other transfections, co-transfection of Nsp1 at a rate 5:1 also knockdown all expression of irrelevant protein expression suggesting at high concentration, Nsp1 was globally repressive for translation which is a known mechanism for the protein (data not shown). Knockout of M2-2 expression was also seen at lower ratios of Nsp1 (data not shown), indicating that the effect on M2-2 by Nsp1 is specific and not simply a result of the overall inhibitory effects of Nsp1.

### Other viral biscistronic systems suggest Nsp1 at lower levels interferes with initiation reinitiation methods of translation

Other viral constructs that required alternative ribosomal translation mechanisms (overlapping ORFs) were also tested against the effects of Nsp1, but only expression of influenza M/BM2 (leaky scanning) (Fig. 4) was affected at 1:1 concentrations. Influenza B M/2 with a upstream 6x his tag and downstream flag expression, when co-transfected with Nsp1, decreased from an average of ∼ eleven cells per field of view to zero to four cells per field of view with no effect on the upstream protein M expression. Nsp1 expression did not appear to knock down Sendai Virus P or C (shunting) expression (Fig.4) or PB1 or F2 (leaky-scanning) expression (Fig.4).

**Figure 4.**
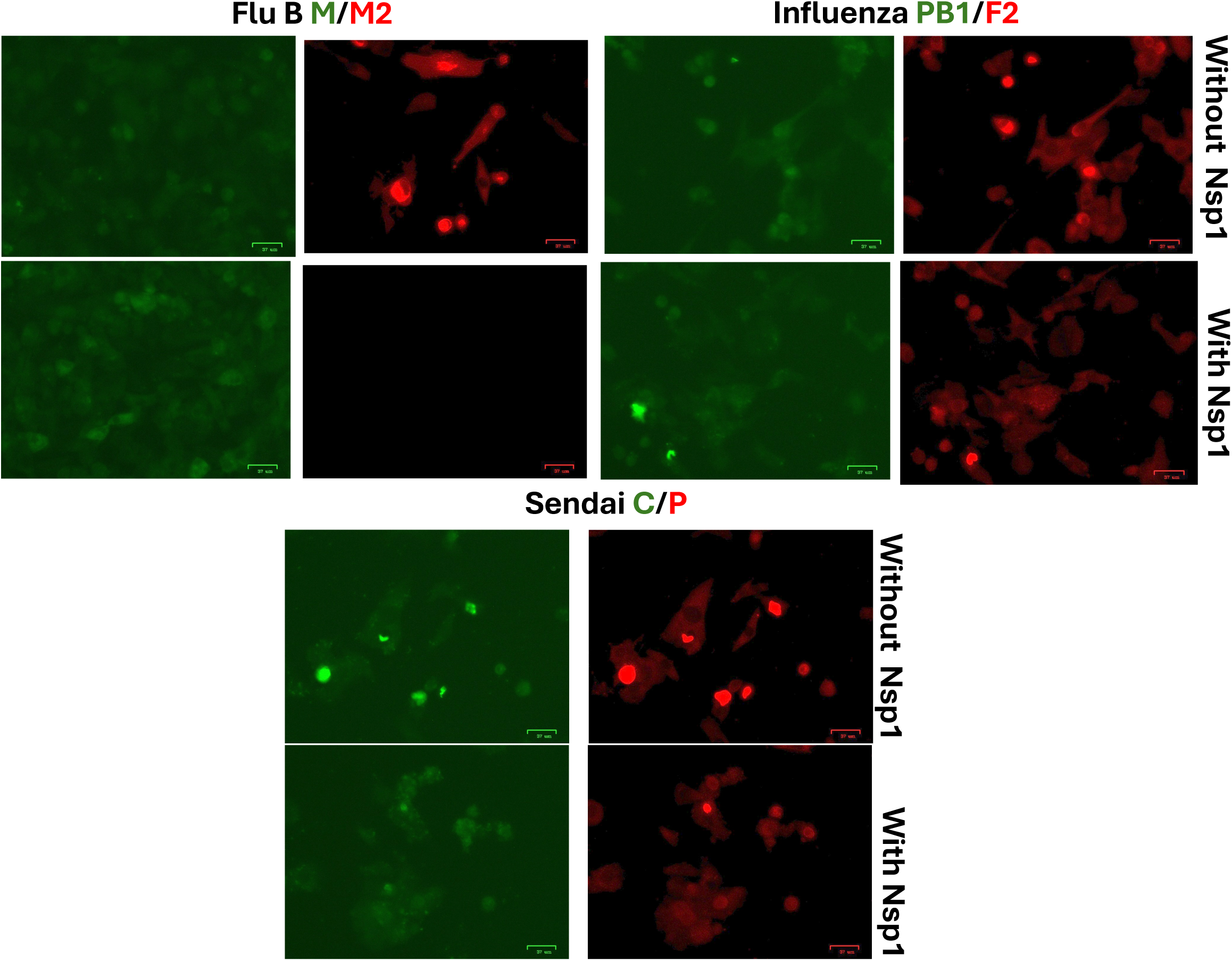
Other bicistronic viral systems suggests that Nsp1 interferes with initiation reinitiation translational methods. We transfected Influenza A PB1/F2, Sendai C/P, Influenza B M1/M2 genes with 6x his or flag tags in each upstream and downstream genes. For some wells Nsp1 was co-transfected at a 1:1 ratio. Shown are **(A)** influenza A PB1/F2 without Nsp1 or **(B)** with Nsp1, **(C)** Sendai C/P without Nsp1 or **(D)** with Nsp1, or **(E)** influenza B M1/2 with Nsp1 or **(F)** without Nsp1. Intracellular staining was similar to Figure 3. Cells in each image were at near confluency when imaged (not shown). n=3 using 3 replicates

### Nsp1 Likely Interferes with 40S Reinitiation in the M2-2 Coding Region

Again, the 40S subunit reads through the M2-1 ORF to the M2-1 termination codon before reinitiating upstream of the M2-2 ATG region and translating M2-2 in conjunction with the other members of the ribosome complex. Prior studies have already shown that the 150-nucleotide region of the M2-1 coding region upstream of the M2-2 ATG start codon is critical for translating the M2-2 protein. We sought to determine where in the M2 mRNA transcript Nsp1 might be competing with the M2-2 ORF for access to the 40S.

The 10ATGM2 construct contained the 10 nucleotides upstream of the M2-2 ATG to allow the ribosome a region to bind the template upstream of the ATG, as constructs that began at the three Gs from the T7 polymerase/kozak/ATG of M2-2 did not express protein for a yet unknown reason despite other genes we tested expressing well when cloned this way (not shown). These constructs were co-transfected with Nsp1 in 1:1 concentrations similar to Figure 3. The 260M2 construct M2-2 expression was knocked out in the presence of Nsp1 (average of 12 cells per field of view reduced to zero to three cells per field of view) (Fig 5A-B), whereas the expression of the 10ATGM2 construct was not knocked down in the presence of Nsp1as. Our 150M2 is not shown but followed a similar trend as 260M2. Thus, Nsp1 likely affects a 40S recruitment element of the M2-2 ORF downstream of the M2-2 ATG site likely in the secondary RNA structure identified by Easton et al (Gould and Easton, 2005).

**Figure 5.**
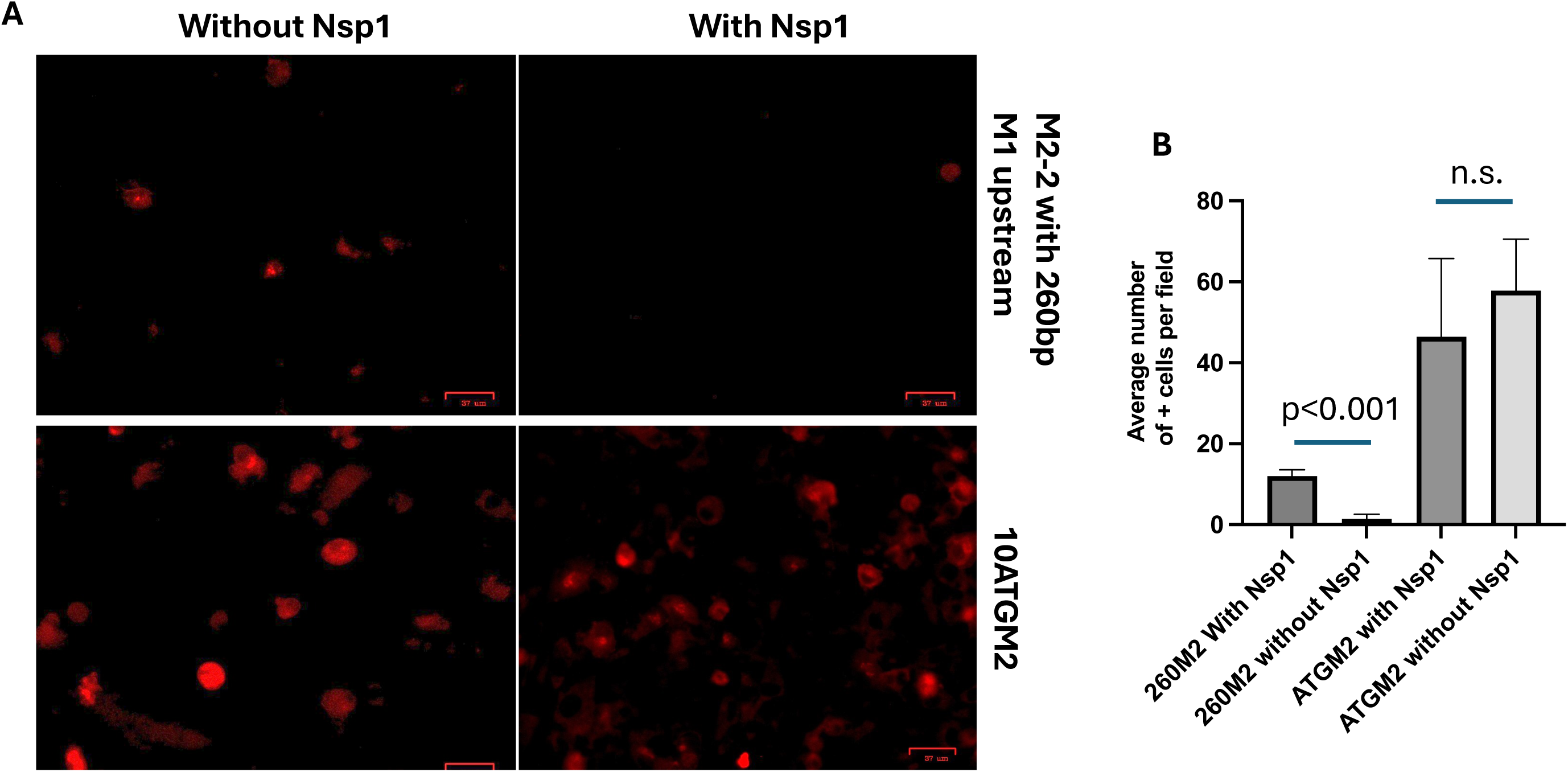
Nsp1 does interfere with M2-2 expression when dependent on its upstream reinitiation site. We transfected ATGM2-2 transcript **(A)** with or **(B)** without Nsp1 and stained for flag intracellualy. Next, we transfected M2-2 with the 150nucleotide upstream portion of M2-1 **(C)** without Nsp1 or **(D)** with it or the 260 nucleotide upstream portion of M2-1 **(E)** with Nsp1 or **(F)** without it. Cells in each image were at near confluency when imaged (not shown). n=3 using 3 replicates

### Nsp1 mutations in the RNA binding sites disrupt its ability to interfere with M2-2 production

Nsp1 mutations at the active 40S/mRNA binding sites (K or R amino acids, locations and mutations in methods) were made and transfections with M-2 done again. We found, as compared to wildtype Nsp1, mutations in either site disrupted the ability of Nsp1 to interfere with M2-2 translation from the bicistronic M2 transcript (Fig. 6A-D) as compared to unmutated Nsp1.

**Figure 6.**
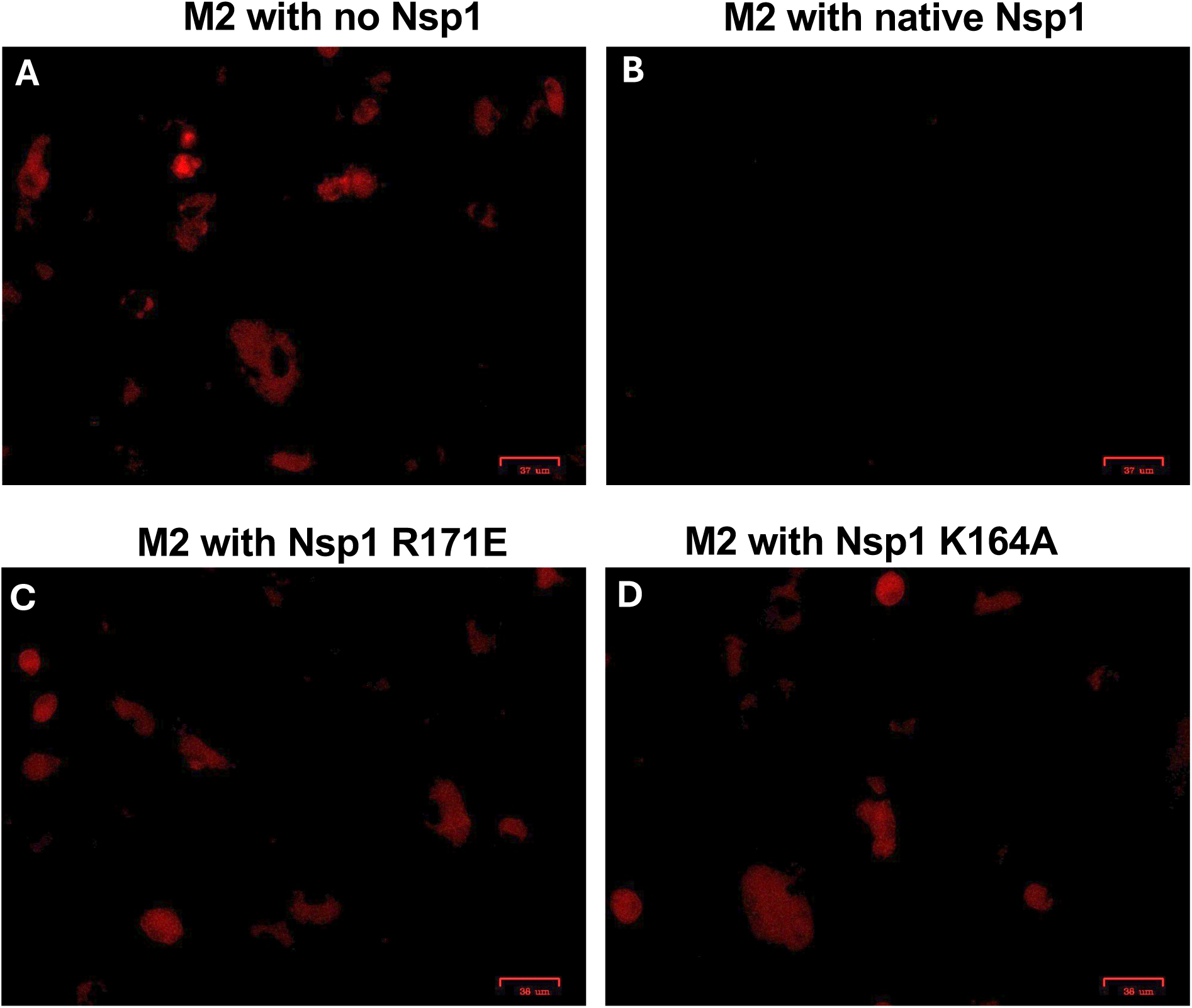
Mutations in Nsp1’s ability to bind to the translational machinery inhibits its block on M2-2 expression. We transfected our M2-1/2 plasmid similar to figure 3 again but using Nsp1 with a mutation in the two regions that allow it to associate with the translational machinery and likely bind mRNA. Shown are **(A)** M2-1/2 expression without Nsp1, **(B)** M2-1/2 expression unmutated Nsp1 **(D)** expression of both proteins with Nsp1 carrying a mutation in its R171E site, and **(D)** carrying a mutation in its K164A site. Cells in each image were at near confluency when imaged (not shown). n=3 using 3 replicates

### Nsp1 Expression Reduces RSV Expression in a Co-transfection-infection Model

BSRT7 cells were transfected with Nsp1 and infected with RSV (mCherry) simultaneously or separately (BHK cells are productively infected by RSV). When cells were transfected with Nsp1 prior to RSV infection, the number of cells expressing the mCherry protein and therefore being infected with RSV were vastly reduced almost to the point of non-detection (Fig. 7A, D, E). In contrast, the RSV infection that was cotransfected with pUC19 with a T7 promoter added in a similar manner had ∼90% of cells/field of view expressing mCherry (Fig. 7B, E). When Nsp1 was transfected into cells that had been already infected with RSV, the level of positive RSV infection was decreased, and while 90% of cells/field of view still expressed mCherry, the average level of mCherry expression was reduced by ∼20% ( average pixel saturation across representive field of view). The transfection of pUC19 did not affect RSV expression of mCherry (Fig.7C). In an examination of effects prior to 24hrs, we found a similar effect on mCherry (RSV replication) expression at 6 hrs if Nsp1 was present.

**Figure 7.**
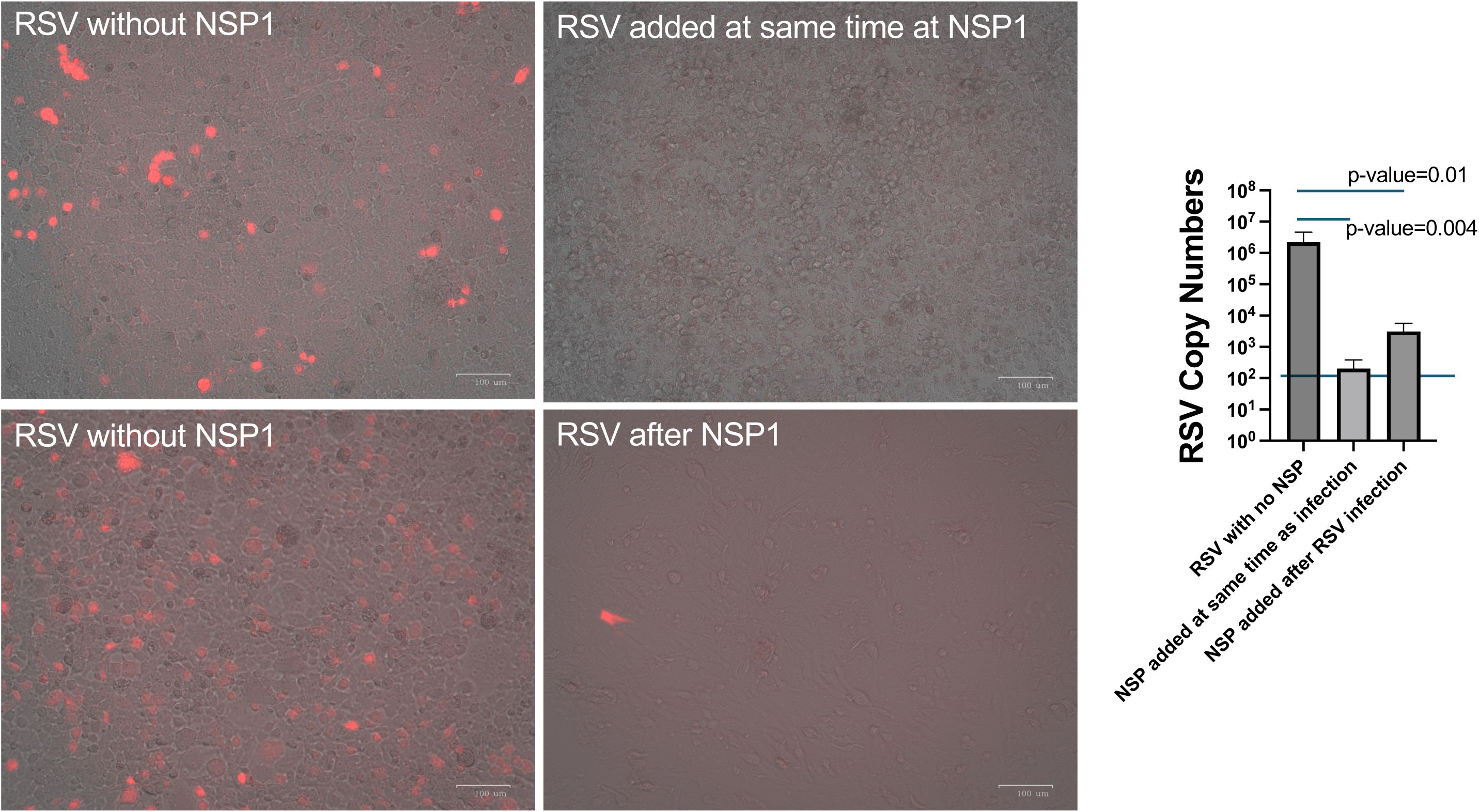
Nsp1 transfection at the same time as infection with a fluorescent reporter RSV confirms it inhibits viral replication. Nsp1 was transfected into BSRT7 cells at the same time as an MOI1 infection of RSV that carries an mCherry reporter behinds it’s NS2 gene. Shown are **(A)** RSV without Nsp1 or **(B)** with it. **(C)** qRT-PCR results for detection of the RSV N gene confirmed our results. Line is our limit of detection in this assay (noise if CTs after 35 from negatives). Images are far red images overlaid with a brightfield image from the same locations. n=3 using 3 replicates

## Discussion

Both RSV and SARS-CoV-2 are respiratory viruses that continue to have significant impacts on human health. With the further relaxation of COVID control measures, typical respiratory viruses should begin to re-circulate at their average numbers. Given SARS-CoV-2’s mutation rate, continuous circulation of this virus is also probable for the long term or at least until better strain matched and vaccine compliance/availability begin to take effect. Thus, while dual infections with SARS-CoV-2 and viruses such as RSV or influenza have not been wide-spread, they likely will be in the future.

Here, coinfection of RSV with SARS-CoV-2 limited the former’s replication within permissive cells which correlated with an absence of M2-2 expression in the presence of Nsp1. Disruption of the RSV M2-2 protein expression by the SARS-CoV-2 Nsp1 protein expression may explain why we observed a limited replication, and not a total elimination of RSV. M2-2 accumulation during the course of RSV infection is a crucial time marker for the RSV lifecycle as accumulation leads to a conformational change in the polymerase complex (Cheng et al., 2005; Sutto-Ortiz et al., 2023). This conformational change in the polymerase structure causes the polymerase complex to switch from a primarily transcriptive state to a primarily replicative one. Without a switch to a replicative state, the virus likely struggles to produce full length antigenome and thus new genomic strands stalling the infection as new empty progeny virions do not have genomes to incorporate and only a small percentage of the progeny are able to obtain a genome and bud out of the cell thereby attenuating infection (Bermingham and Collins, 1999; Blanchard et al., 2020). Inhibition of expression in both the 150M2 and 10ATGM2 constructs by Nsp1 indicates that Nsp1 is likely affecting M2-2’s coding sequence (indirectly through the ribosome or host endonucleases) rather than interacting with any regulatory elements upstream in the 150 region. The inhibition of the 10 nucleotide substitution construct indicates that this is likely some distance down-stream of the ATG sequence. If Nsp1 is directly binding or leading to cleavage of the M2 mRNA in such a manner that prevents M2-2 expression, then ribosomal profiling experiments designed to map the exact regions Nsp1 affects, or RNA protection assays would also provide tantalizing information to design inhibitory molecules which could act as RSV therapies by attenuating M2-2 expression and thus RSV infection.

While we concentrated on the M2-1/M2-2 switch, Nsp1 could affect other parts of the RSV replication cycle as well. Alternatively, Nsp1 effects on host mRNA translation could also rob RSV of critical host factors needed in its replication. Future studies would entail using our reverse genetic system to make reporters in the RSV M2 genome to allow us to determine if a similar limitation of the M2-2 switch is occurring during co-infection. Ongoing studies using Nsp1 transfections with RSV infection suggest that Nsp1 is largely responsible for the viral attenuation of RSV in the presence of SARS-CoV-2. Differences in RSV strains could also be impacted differently by Nsp1. For instance, we used the RSVa 2001 clinical strain, which contains two in-frame ATG start sites separated by only nine nucleotides in the M2-2 protein-coding region, while the more frequently used lab strain of RSV (RSVA2) has three in-frame ATG sites. In another study using SARS-CoV-2 with RSVA2 strain in vitro, there was a marked reduction in the former (Dee et al., 2023). These further evidence that the effects of each virus on each other might be strain specific.

While influenza A and B were significantly reduced by SARS CoV-2 infections as were most other respiratory viral infections, new studies using influenza A and SARS CoV-2 demonstrate that simultaneous infections can increase the level of infection for both viruses in cells or permissive animals (Kinoshita et al., 2021). We did see potentially higher PB1 upstream expression in the presence of Nsp1 but that needs to be further explored. We are unaware of studies looking at synergy between influenza B or parainfluenzas with SARS CoV-2. Another study continues to show low co-infection rates between influenza B strains and SARS CoV-2 (Yan et al., 2023) but interference between these two viruses or synergy would be interesting to explore in more depth.

We demonstrate a SARS-CoV-2 effect on RSV replication during a co-infection in vitro model. The effect appeared to be targeted to the M2-2 downstream protein on the M2-1/M2-2 bicistronic mRNA strand. While we would think using Nsp1 as a therapeutic for RSV infections as Nsp1 could disrupt the replication of the RSV, the potentially deleterious effects of Nsp1 on host protein translation could make using the protein as a therapy difficult. Instead, further investigation into the exact mechanism of action of Nsp1 on M2-2 protein expression may reveal more specific targets allowing for a therapeutic to be developed that would not target global mRNA translation the way Nsp1 does.

## Abbreviations

Hr: Hour

## Acknowledgements

Supported in part by a research grant to DV from Investigator-Initiated Studies Program of Merck Sharp & Dohme LLC. The opinions expressed in this paper are those of the authors and do not necessarily represent those of Merck Sharp & Dohme LLC.

## Figure Legend

**Supplemental Figure 1.**
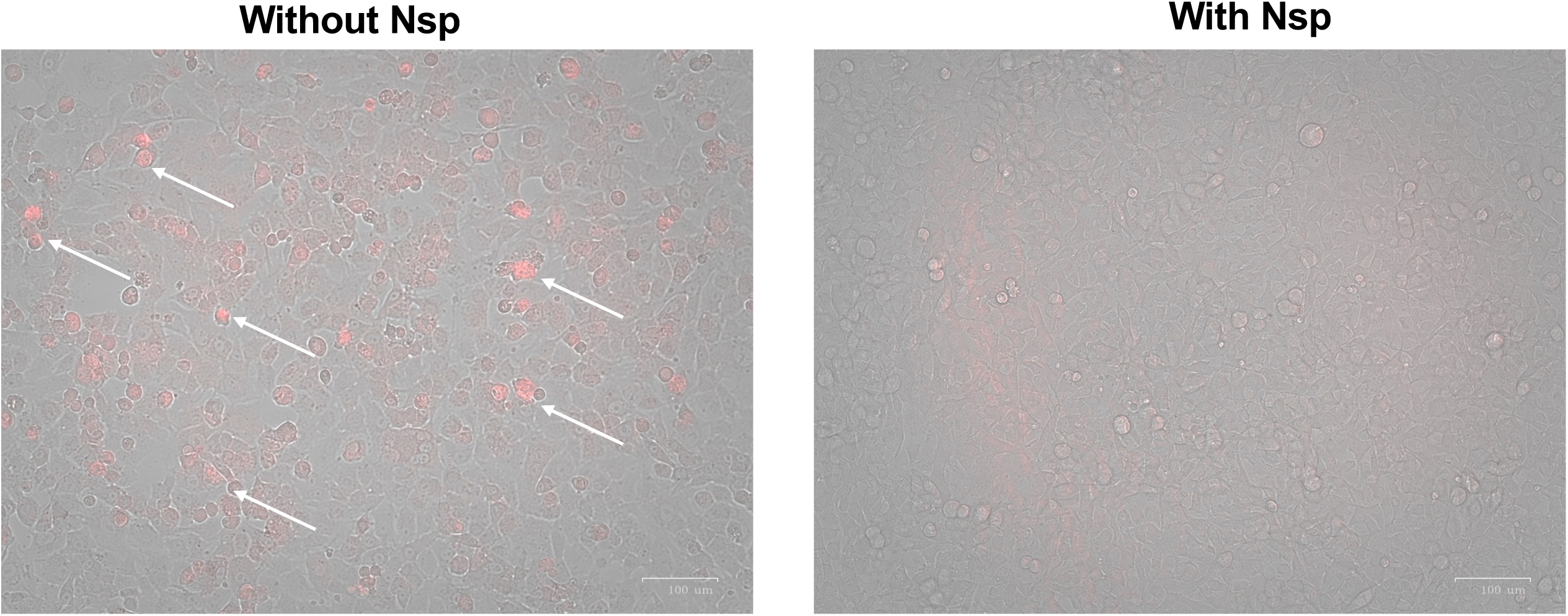
Nsp1 blocks virus very early into the infection. We infected BSRT7 cells with fluorescent reporter virus at the same time transfected Nsp1. We imaged for mCherry expression by 6 hrs post-infection.

